# Multicopper oxidase mediated single-carbon insertion for skeletal remodeling

**DOI:** 10.64898/2026.03.28.714988

**Authors:** Bo Jiang, Binbin Chen, Hongxun Gao, Jiaxin Huang, Xiao Liu, Mingjie Ma, Yajie Wang

## Abstract

Modern drug discovery demands efficient strategies for generating structurally diverse compound libraries. Skeletal editing—a transformative paradigm enabling precise atom-level modifications within molecular frameworks, offers a sustainable alternative to traditional synthetic routes. While carbene insertion-mediated approaches have dominated single-carbon insertion strategies, current methodologies are limited by their reliance on hazardous, unstable carbene precursors and harsh reaction conditions. Herein, we report a multicopper oxidase (MCO)-catalyzed skeletal editing that enables the direct, one-step transformation of phenolic and indole derivatives into functionalized tropones and quinoline analogues through exogenous single-carbon insertion. This platform employs stable and safe nitroalkanes as carbon sources and O_2_ as the sole terminal oxidant. It accommodates a broad substrate scope and yields products with superior antibacterial activity against to multidrug-resistant strains relative to their parent compounds. This work introduces the first biocatalytic platform for exogenous single-carbon insertion skeletal editing. This sustainable and scalable strategy overcomes key limitations of synthetic approaches, offering efficient skeletal remolding and rapid expansion of bioactive compound libraries.

**Graphic Abstract:** 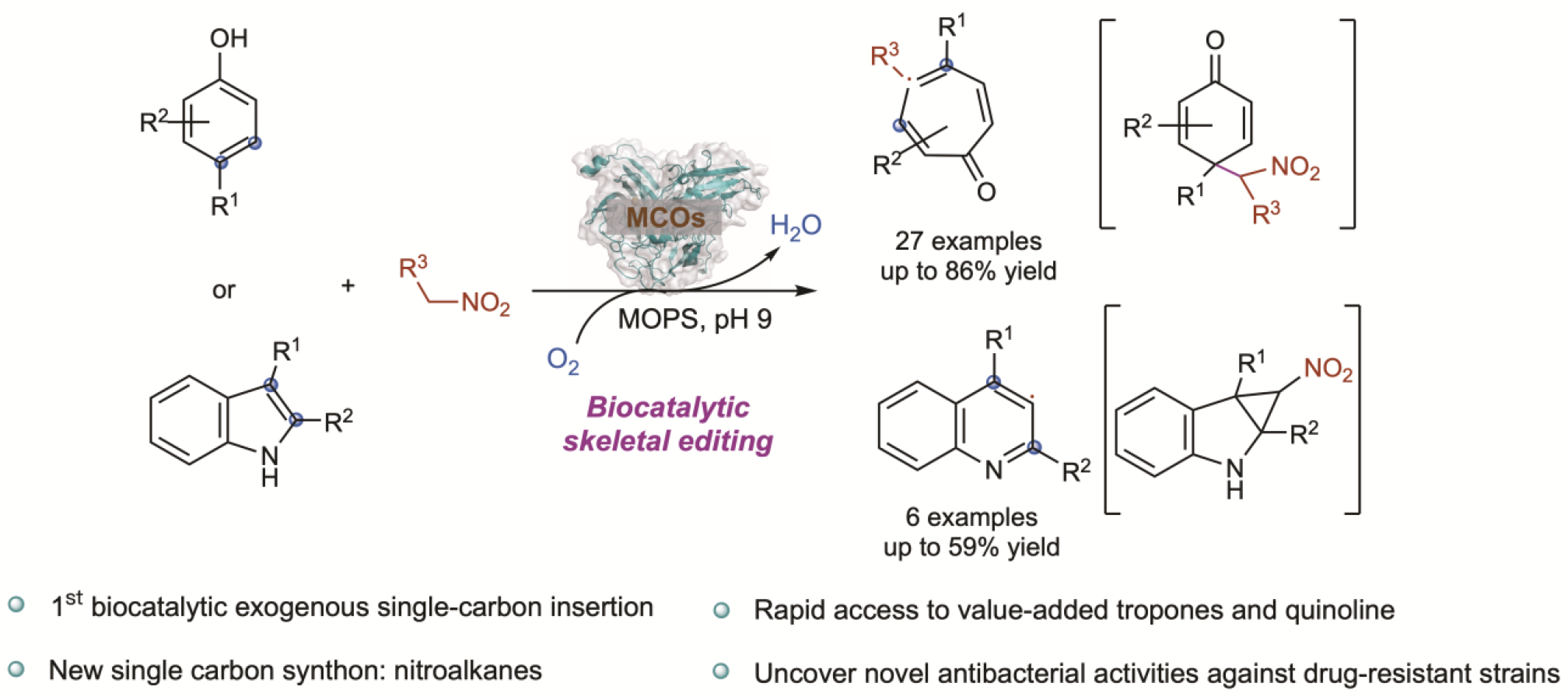

## Main

A key demand in contemporary drug discovery is the development of efficient methods to produce compound libraries with rich structural diversity^1-3^. Skeletal editing—an emerging approach in synthetic chemistry—addresses this demand by enabling precise atom-level modifications (insertion, deletion, or exchange) within molecular frameworks^4,5^. This strategy retains core structural elements while fine-tuning physicochemical and bioactive properties, offering a more efficient and sustainable alternative to traditional multistep *de novo* synthesis. Notably, single-atom insertion into cyclic systems is particularly promising for enabling rapid access to pharmaceutically relevant expanded-ring scaffolds (Fig. 1a)^6-14^.

**Fig. 1.**
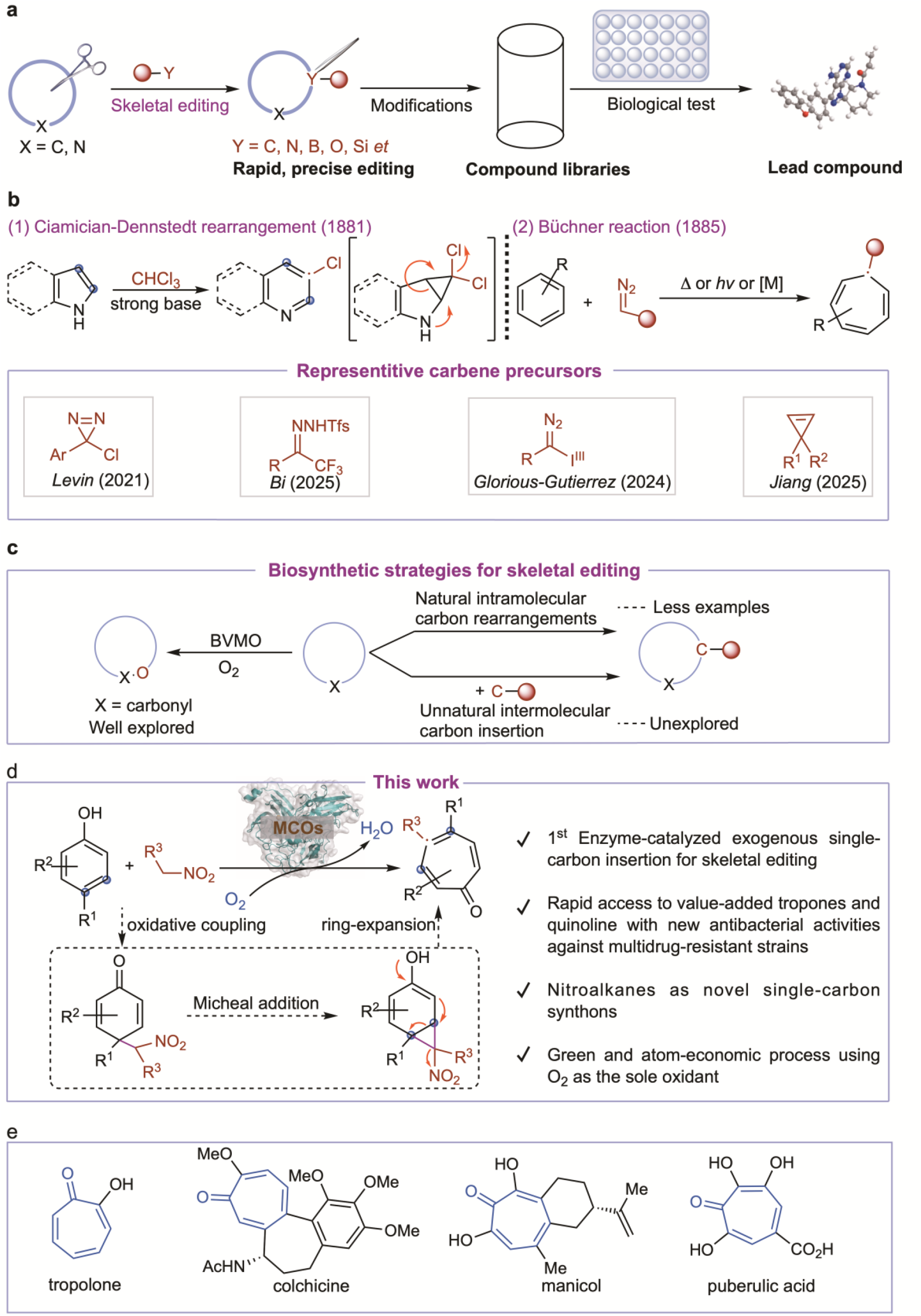
Skeletal editing of cyclic systems via single-atom insertion. **a**, Skeletal editing as an indispensable core technology for constructing diversified drug screening libraries **b**, Classical skeletal editing strategies via single-carbon insertion. **c**, Biosynthetic strategies for skeletal editing. **d**, This work: Multicopper oxidase (MCO)-mediated novel skeletal remodeling strategy for intermolecular single-carbon insertion. **e**, Representative examples of bioactive natural products containing the tropone scaffold.

The molecular editing of cyclic architectures through single-carbon insertion traces its origin to transformations such as the Ciamician-Dennstedt^15^ rearrangement and the Büchner^16^ reaction (Fig. 1b-1,2). These methods utilize carbene intermediates to incorporate a single carbon atom into pyrroles and benzene rings. Recent advancements have significantly broadened this strategy through the development of engineered carbene precursors, including aryl chlorodiazirines^17^, trifluoromethyl *N*-triflyl hydrazones^18^, and α-iodonium diazo compounds^19^, etc.^20,21^, enabling efficient carbon insertion into nitrogen-containing aromatic heterocycles and indene. Notably, a recent rhodium/boron catalytic system achieved the first carbene-mediated single-carbon insertion into phenol derivatives^22^. Despite these advancements in constructing molecular scaffold diversification, the practical application of carbene-mediated insertion is hampered by several limitations. The requirement for pre-preparation of unstable and potential explosive carbene precursors, coupled with harsh reaction conditions, frequently results in low yields, cumbersome operations, and substantial synthetic costs^23,24^. Moreover, the one-step expansion of six-membered aromatic rings to their seven-membered counterparts pose a significant synthetic challenge^25^. Currently, the predominant method remains the Büchner reaction^26^, which relies on non-sustainable and environmentally hazardous stoichiometric oxidants, limiting its practicality and scalability. These constrains have severely restricted the broader implementation of this skeletal editing strategy. Consequently, the development of readily accessible, stable, and safe single-carbon insertion reagents, coupled with greener oxidative process, is urgently needed to conventional carbene precursors and advance the skeletal editing of six-to seven-membered aromatic rings.

Over the past decade, significant advances have been made in engineering enzymes via directed evolution to facilitate abiotic transformations, bridging the gap between chemical catalysis and biocatalysis^27-31^. Notable examples include cytochrome P450-mediated carbene and nitrene transfer reactions^32^, non-heme iron enzyme-catalyzed olefin difunctionalization^33^, and visible light-driven radical transformations facilitated by nicotinamide-^34^, flavin-^35^, thiamine-^36^, and pyridoxal-5’-phosphate^37^-dependent enzymes. Although biosynthetic strategies provide valuable platforms for executing challenging chemical transformations, research on molecular skeletal editing remains limited. Current efforts predominantly focus on Baeyer-Villiger monooxygenase (BVMO)-catalyzed single-oxygen insertion into cyclic frameworks^38^. In contrast, strategies for single-carbon insertion remain underdeveloped, with only isolated reports of biotic ring expansions via intramolecular carbon rearrangements multi-enzyme catalyzed cascade reactions^39-41^. To date, biocatalytic intermolecular single-carbon insertion for skeletal remodeling remains entirely unexplored, representing a critical gap in programmable skeletal editing technologies (Fig. 1c).

Herein, we report an innovative skeletal editing strategy catalyzed by multicopper oxidase (MCO) to convert abundant phenol derivatives into functionalized tropones via exogenous single-carbon insertion in one step (Fig. 1d). This transformation employs cost-effective, safe, and stable nitroalkanes as novel single-carbon synthons and O_2_ as the sole oxidant, circumvents the limitations of traditional carbene-based insertion methods, which have dominated advancements in single-carbon insertion for skeletal editing over the past decade. Mechanistically, this transformation diverges from classical MCO-catalyzed pathways of phenol derivatives self-coupling or quinoid intermediates formation ^42^. Instead, it proceeds through a radical oxidative addition between phenols and nitroalkanes, generating dienone. Subsequently, a spontaneous tandem sequence of Michael addition and ring-expansion proceeds through a norcaradiene intermediate, yielding diverse tropone derivatives, which are versatile synthetic motifs^43^ and core scaffold of numerous bioactive compounds^44-46^ (Fig. 1e). Notably, this work establishes the first biocatalytic platform for skeletal editing via exogenous single-carbon insertion, enabling ring expansion of phenol and indole systems to directly yield functionalized tropones and quinolines, respectively. Several of these synthesized compounds exhibited novel antibacterial activity against multidrug-resistant strains compared to their starting substrates. Overall, this transformation establishes an unprecedented paradigm in synthetic and enzymatic catalysis, significantly expanding the frontiers of skeletal remodeling.

## Results

### Reaction design and optimization

Inspired by natural biosynthetic pathways employing multicopper oxidase (MCO) for oxidative coupling of phenol derivatives^42^, we hypothesize that norcaradiene^47^, a crucial intermediate for ring-expansion via single-carbon insertion, could be generated through an MCO-catalyzed oxidative coupling between phenol derivatives and activated alkylation substrates, followed by a spontaneous intramolecular Michael addition. To test this hypothesis, we first investigated the reaction between 3,5-dichloro-*p*-toluene-4-sulfonamido phenol (**1a**) and various alkylation substrates, including nitromethane (**2a**), acetophenone (**2b**), methyl *p*-tolyl sulphoxide (**2c**), and methyl *p*-tolyl sulfone (**2d**), catalyzed by MCOs from *Escherichia coli* K-12 (Ec-Lac). The desired ring-expansion product was obtained only when **2a** was employed (Fig 2a, entry 1). To improve reactivity, we screened additional MCOs identified in our previous study^48^. Although most did not enhance conversion, **BacFre** from *Bacillus freudenreichii*, afforded a modest increase in yield (Fig 2a, entry 8). We subsequently performed site-saturation mutagenesis on residues within 6 Å of T1 copper active site. After two rounds of directed evolution, the variant **BacFre-K474F/I500L** exhibited a threefold higher yield than the wild type, reaching 30.3% (Fig. 2b). Through systematic optimization of reaction parameters, including nitromethane (**2a**) stoichiometry, whole-cell catalyst concentration (OD_600_ = 10) (Fig. 2c), buffer system, and oxidants, we achieved an 88% yield of tropone **3a** using MOPS buffer (pH 9) and 30 equivalents of **2a** (Fig. 2d). A similar result was obtained using purified enzyme **BacFre-K474F/I500L**. Control reactions employing either the PET28a empty vector or CuSO_4_ produced only trace amount of the product, confirming that the catalytic oxidative ring-expansion is specifically mediated by the evolved MCO variant **BacFre-K474F/I500L** (Fig. 2e).

**Fig. 2.**
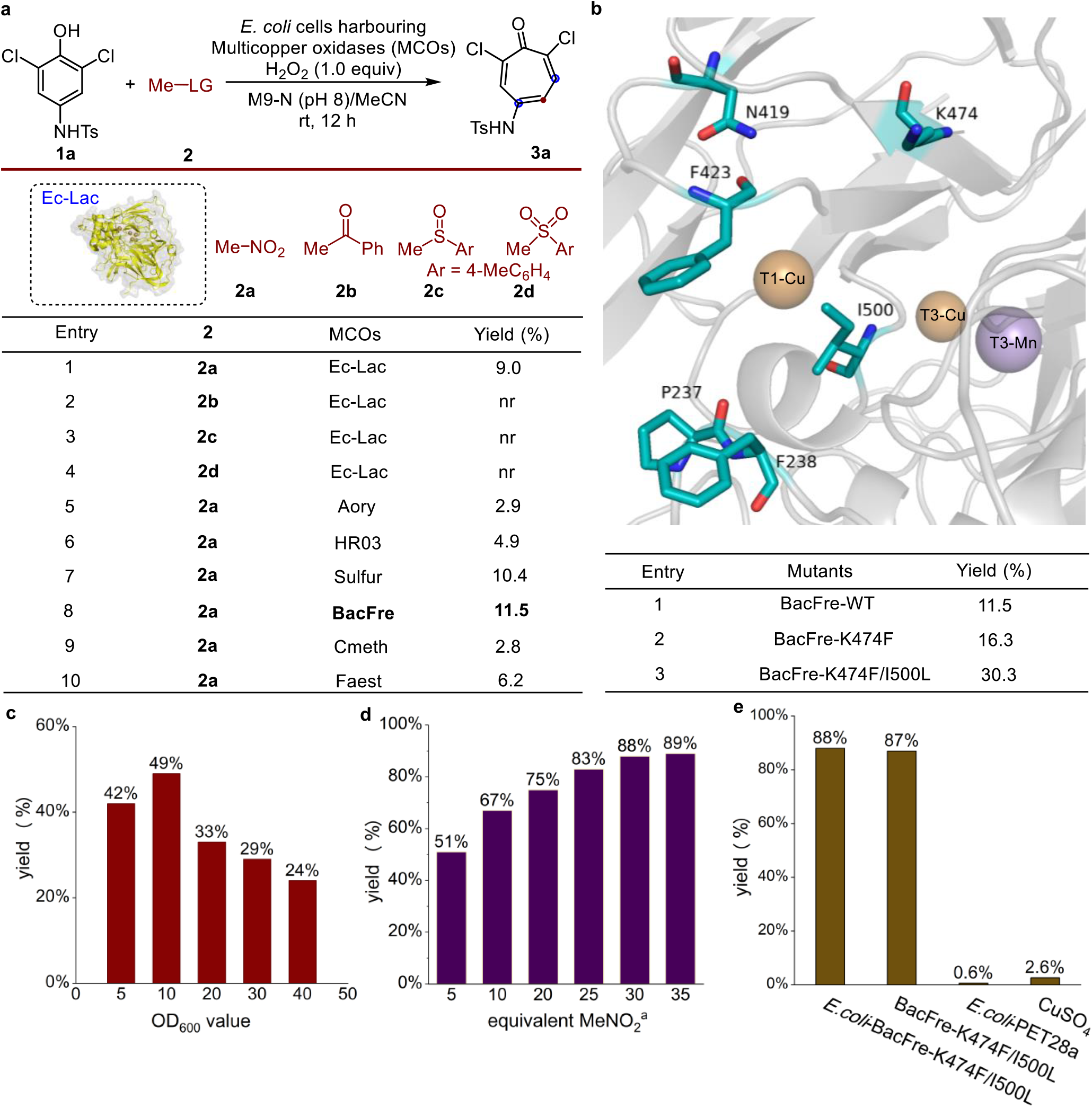
Development of MCO-mediated skeletal editing. **a**, Screening of alkylation reagents and MCOs for skeletal editing via single-carbon insertion. **b**, Representative active site of **BacFre** and engineering results (yield improvement of key variants; see Supplementary Table 2 for full mutagenesis data). **c**, Effect of whole-cell catalyst density (OD_600_ value) on product yield. **d**, Effective of nitromethane stoichiometry on yield (for screening of buffer system and oxidants, see Supplementary Table 5 and 6). **e**, Control experiments confirming enzyme-dependent catalysis. Reaction conditions: **1a** (0.004 mmol, 40 µL of 0.1 M solution in MeCN), alkylation reagents **2** (0.02 mmol, 10 µL of 2.0 M solution in MeCN), H_2_O_2_ (0.004 mmol, 1 µL), suspensions of *E. coli* whole cells expressing MCOs (OD_600_ = 20) in M9-N buffer (350 µL, 50 mM, pH 8). Reactions were performed at room temperature under air for 12 h. Yields were determined by reversed-phase liquid chromatography using benzophenone as an internal standard. ^a^*E. coli* whole cells expressing **BacFre-K474F/I500L** (OD_600_ = 10) in MOPS buffer (50 mM, pH 9), without H_2_O_2_. LG: leaving group.

### Substrate scope investigations

With the optimal conditions established, we evaluated the substrate scope of **BacFre-K474F/I500L**-catalyzed skeletal editing via single-carbon insertion (Fig. 3). Initial performing reactions with diverse sulfamido phenol substrates with nitromethane (**2a**), revealed broad functional group tolerance, including various aryl and heteroaryl sulfones (**1a**–**1m**), affording tropones **3a**–**3m** in moderate to excellent yields (32–86%). In addition, the reaction also accommodated alkyl-substituted derivatives bearing *N*-mesyl (**1n**) or *N*-cyclopropyl sulfonyl (**1o**) group, affording products **3n** and **3o** in 31% and 38% yield, respectively. Moreover, evaluation of phenol substitution patterns demonstrated successful conversion of 3,5-dibromo-(**1p**), 3-chloro-*p*-toluene-4-sulfonamido phenol (**1q**) and unsubstituted *p*-toluene-4-sulfonamido phenol (**1r**) to the corresponding tropones (**3p**–**3r**) with 20–68% yield. In contrast, 2,5-dichloro-*p*-toluene-4-sulfonamido phenol (**1s**) exhibited low reactivity, yielding only 7% of **3s**. Notably, the 3-phenyl-*p*-toluene-4-sulfonamido phenol (**1t**) afforded both the target tropone (**3t**) and the *p*-hydroxybenzaldehyde derivative **4a** as a byproduct. Further investigations showed that substrates with electron-donating groups at the 3-position (**1u**–**1w**) preferentially formed *p*-hydroxybenzaldehyde derivatives **4a**–**4d**. We propose that these byproducts arise from ring contraction and decomposition of the corresponding tropones when electron-donating groups are present (see Supplementary Fig. 19). Meanwhile, 2-substituted *p*-toluene-4-sulfonamido phenol (**1x** and **1y**) failed to yield the ring-expansion product and instead formed dimeric species **5a** and **5b** exclusively. We attribute this divergent reactivity to the heightened propensity of 2-substituted phenols to undergo oxidative dimerization, a pathway favored by the reduced steric hindrance at the 3-position, which outcompetes the desired reaction with nitromethane. In addition, the reaction demonstrated good scalability, with product **3a** was obtained in comparable yield on 1.0 mmol scale.

**Fig. 3.**
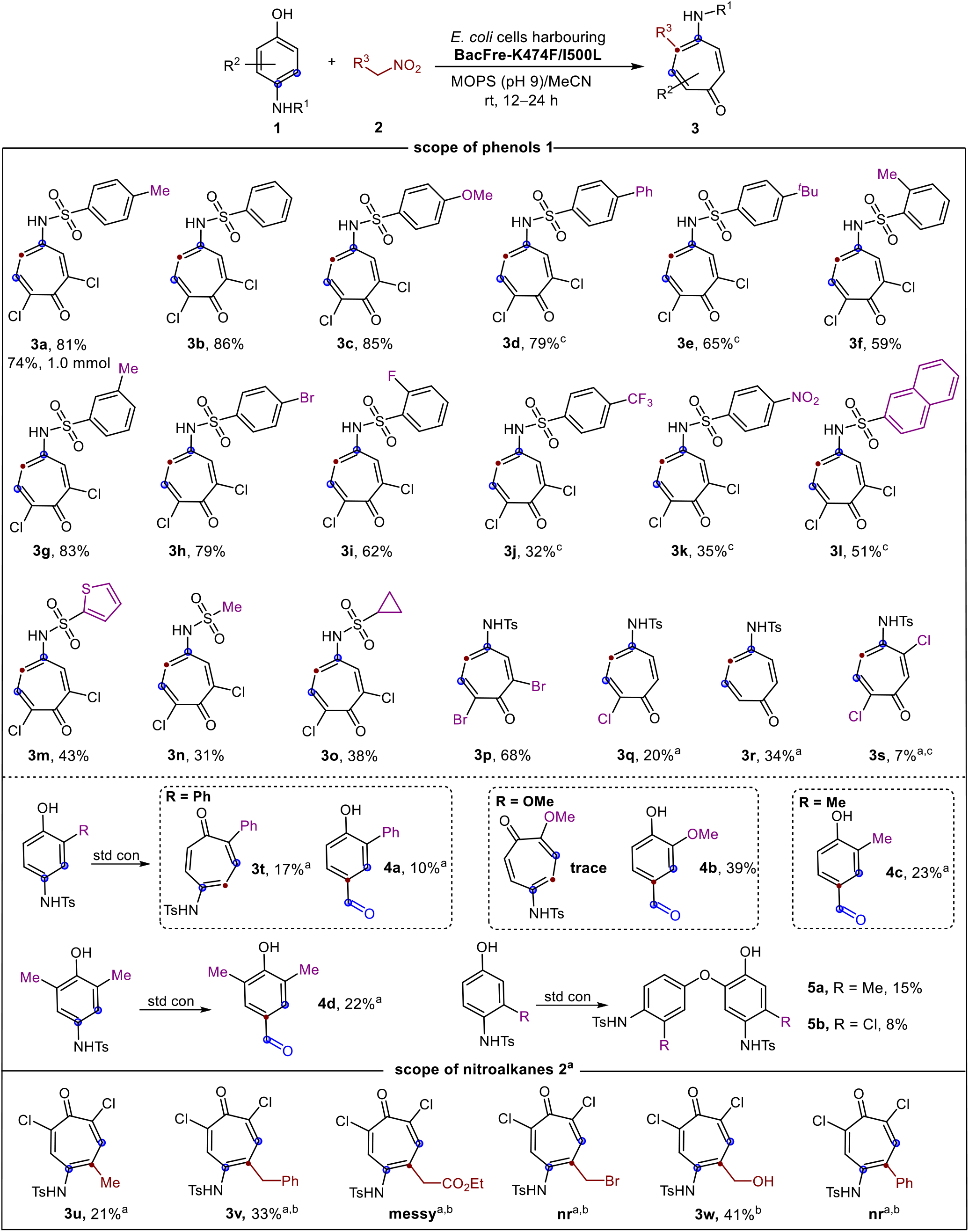
Substrate scope of enzymatic skeletal editing via single-carbon insertion. Unless otherwise noted, reactions were carried out with **1** (0.2 mmol) and **2** (6 mmol) in MeCN (2 mL), suspensions of *E. coli* expressing **BacFre-K474F/I500L** (OD_600_ = 10) in MOPS buffer (18 mL, 50 mM, pH 9). Reactions were conducted at room temperature under air for 12 h. Isolated yield. ^a^*E*.*coli* expressing **BacFre-K474F/I500L** (OD_600_ = 20). ^b^Nitroalkanes **2** (2 mmol). cReaction for 24 h. The absolute configuration of **3a** and **4d** was determined by X-ray crystallographic analysis. All other products were assigned by analogy. Std con: standard condition.

The direct utilization of diverse nitroalkanes (**2**) bearing various functional groups would significantly expand the scope of this skeletal editing strategy, enabling access to multi-substituted tropones. As illustrated in Fig. 3, nitroethane (**2e**) proved applicable, albeit affording product **3u** in moderate yield (21%). Several functionalized nitroalkanes were evaluated, including benzyl, ester-, bromo-, hydroxyl-, and phenyl-substituted derivatives (**2f**–**2j**). While (2-nitroethyl) benzene (**2f**) and 2-nitroethan-1-ol (**2i**) delivered tropones **3v** and **3w** in moderate to good yield (33% and 41%), other substituted nitroalkenes did not yield the desired tropone products.

Except phenols, this strategy was successfully extended to the single-carbon insertion and ring expansion of indole substrates **6a–6f**, yielding quinoline products **7a–7f** in 4%**–**59% yields (Fig. 4). Quinoline represents a privileged organic scaffold prevalent in many natural products and pharmaceuticals, such as quinolone antibiotics, the antiviral agent saquinavir, and the antimalarial drug chloroquine^49^. Finally, our catalytic system was successfully used to realize direct single-carbon mediated ring expansion of several bioactive compounds (Fig. 4). Antiangiogenic agents (**8**)^50^, mTORC1 inhibitors (**9**)^51^, Z α1-antitrypsin antagonists (**10**)^52^, and PI4KIIIβ inhibitors (**11**)^53^ were converted to their corresponding seven-membered ring products (**12**–**15**) under standard conditions using **2a**, yielding fair to moderate isolated yields (11–36%).

**Fig. 4.**
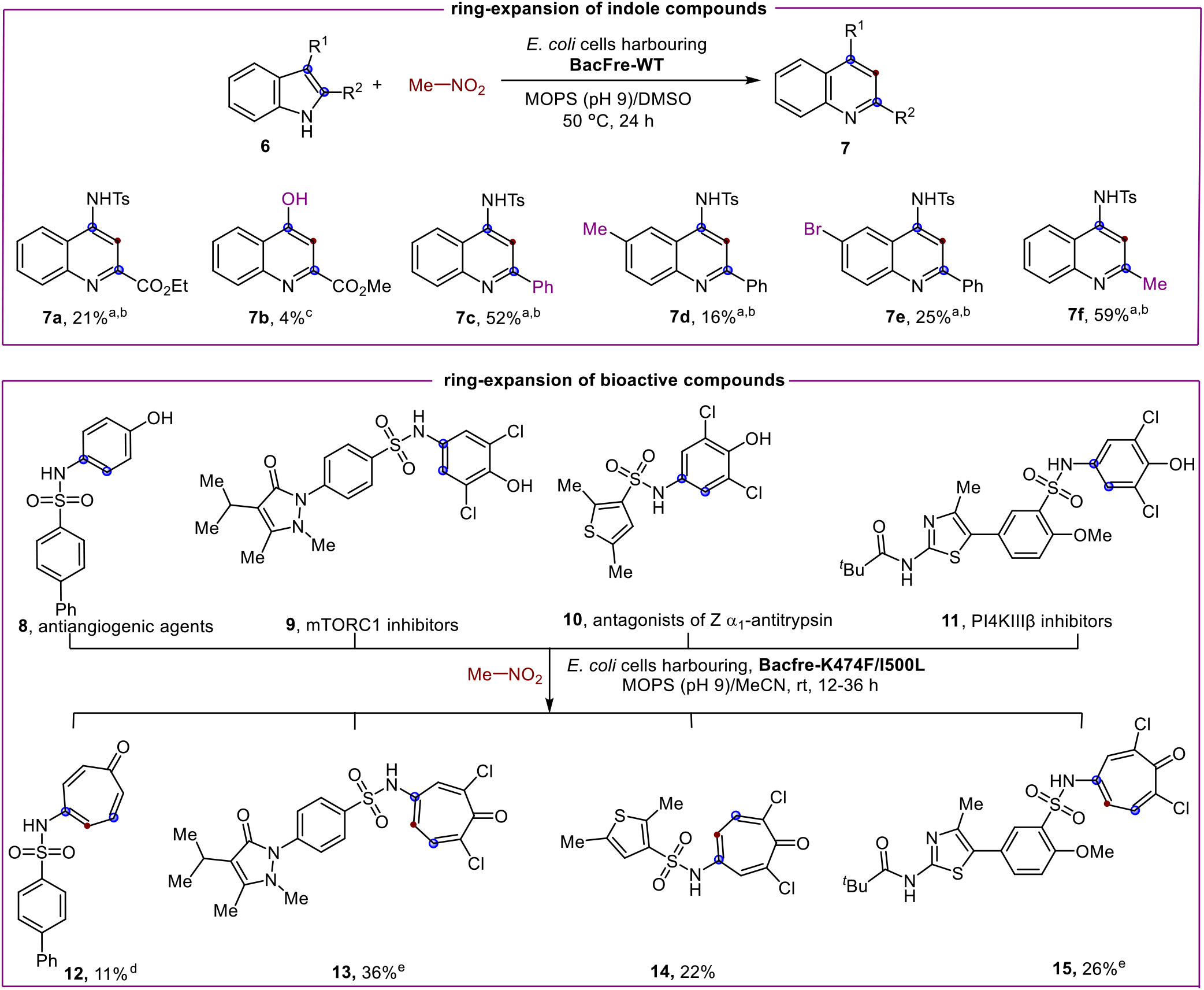
Application of MCO-catalyzed skeletal editing strategies to indole substrates and bioactive molecules. Unless otherwise noted, reactions were carried out with indole substrates or bioactive molecules (0.2 mmol), **2a** (6 mmol) in MeCN or DMSO (2 mL), and suspensions of *E. coli* expressing **BacFre-K474F/I500L** (OD_600_ = 10) in MOPS buffer (18 mL, 50 mM, pH 9). Reactions were conducted at room temperature under air for 12 h. Isolated yield. ^a^*E*.*coli* expressing **BacFre-WT** (OD_600_ = 30). ^b^Reaction performed at 50 °C. ^c^*E*.*coli* expressing **BacFre-K474F** (OD_600_ = 30) in MOPS buffer (18 mL, 50 mM, pH 5.5). ^d^*E*.*coli* expressing **BacFre-K474F/I500L** (OD_600_ = 20). ^e^Reaction for 36 h.

We also evaluated the antibacterial activity of the tropones products alongside their parent substrate against a panel of multidrug-resistant Gram-positive and Gram-negative bacteria pathogens (Table 1, Supplementary Fig. 20 and 21). Compound **3a** exhibited potent activity against Gram-negative strains *Riemerella anatipestifer* (*RA4049*), whereas its precursor **1a** showed no activity. Similarly, compounds **3p** and **3s** displayed notable potency against both *RA4049* and Gram-positive strains *Streptococcus dysgalactiae* (*T8*). while their corresponding substrate **1p** and **1s** were inactive. Those results demonstrate that this strategy holds transformative potential for medicinal chemistry, providing a powerful tool for late-stage molecular diversification and expanding the accessible chemical space in drug discovery.

**Table 1.** Evaluation antibacterial activity of the tropone products and substrates.

| Strains | potency | 1a | 3a | 1p | 3p | 1s | 3s |
| --- | --- | --- | --- | --- | --- | --- | --- |
| **T8** | IZ (mm) | — | — | — | 8.15±0.91 | — | 7.64±0.40 |
|  | Conc. (mg/mL) | — | — | — | 0.25 | — | 0.10 |
| **RA4049** | IZ (mm) | — | 7.23±0.20 | — | 7.88±0.19 | — | 11.95±0.24 |
|  | Conc. (mg/mL) | — | 0.25 | — | 0.25 | — | 0.10 |

Note: “—” indicates no potency. IZ: inhibition zone, the mean of triplicate vernier caliper measurements.
Conc.: the chemical stock concentration used for the IZ measurement with 6-mm Oxford cup

### Mechanistic studies

A possible mechanism involves MCO-mediated oxidative addition of phenol with nitroalkane to form dienone intermediate **INT3**, followed by a cascaded intramolecular Michael addition to generate the key norcaradiene intermediate (**INT5**). Subsequent ring-expansion rearrangement then affords the tropone products (Supplementary Fig. 5). Several experiments were conducted to elucidate the mechanism of this biocatalytic system. Molecular dynamic (MD) simulations identified key residues involved in electron and proton transfer during the phenol oxidation catalyzed by **BacFre** (Supplementary Fig. 6–10). H503 facilitates electron transfer from the substrate to the T1-Cu center, while C498 shuttles electrons from T1-Cu to the T3-Cu/Mn clusters. Mutation of either residue to alanine abolished the catalytic activity (Fig. 5a). Notably, a significant reduction in activity for the E502A mutant suggests a critical role for E502 as a catalytic base in abstracting a proton from the phenolic hydroxyl group (Fig. 5a). Structure analysis further revealed a salt bridge between K474 and E306, which is disrupted by the K474F mutation (Supplementary Fig. 8). We hypothesize that disruption of this interaction enables E306 to act as a general base, facilitating proton abstraction during substrate oxidation and resulting in enhanced catalytic activity. This proposed role is supported by the complete loss of activity observed in the E306A mutant (Fig. 5a). In addition, the improved yield observed for the I500L mutant is consistent with the elimination of a hyperconjugative electron-donating effect upon Ile-to-Leu substitution. This change reduces the electron cloud density of the oxygen atom, facilitating protein dissociation and thereby enhancing oxidation of the hydroxyl group at the T1-Cu center (Supplementary Fig. 10).

**Fig. 5.**
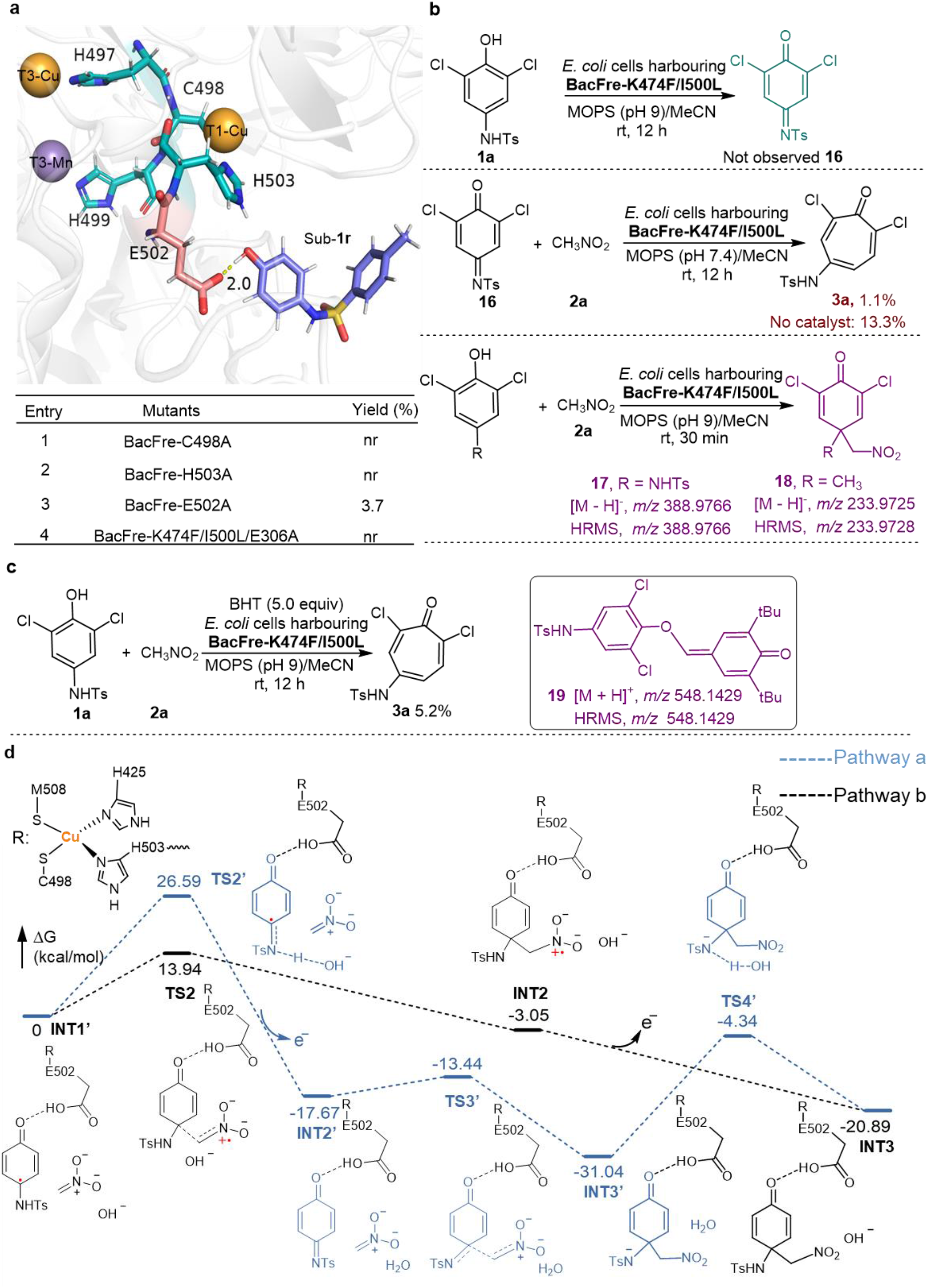
Mechanistic studies. **a**, Molecular dynamic simulations identified key residues of **BacFre. b**, Trapping the critical intermediates of model reaction. **c**, Trapping with BHT (butylated hydroxytoluene) in model reaction. **d**, Calculation of Gibbs free energy diagram (kcal/mol) for **BacFre**-catalyzed oxidative addition from **INT1’** to **INT3**. Unless otherwise noted, reactions were carried out with **1a** (0.004 mmol, 40 µL of 0.1 M solution in MeCN), **2a** (0.12 mmol, 10 µL of 12 M solution in MeCN), suspensions of *E. coli* whole cells expressing **BacFre** variant (OD_600_ = 10) in MOPS buffer (350 µL, 50 mM, pH 9). Reactions were performed at room temperature under air for 12 h. Yields were determined by reversed-phase liquid chromatography using benzophenone as an internal standard.

Two putative pathways were proposed for the formation of dienone intermediate **INT3**. In **Pathway a, INT1** undergoes deprotonation and enzymatic re-oxidation to yield iminoquinone **INT2’**, which then participates in a Henry reaction with nitromethane. Subsequent proton transfer to the nitrogen atom leads to the formation of dienone intermediate **INT3** (Supplementary Fig. 5-**Pathway a**). In contrast, **Pathway b** entails a direct radical oxidative coupling between the carbon-centered radical (**INT1**) and nitromethane, generating a nitro radical intermediate^54^ (**INT2**) that is subsequently oxidized to form dienone intermediate **INT3** (Supplementary Fig. 5-**Pathway b**). Systematic mechanistic investigations were conducted to delineate the operative pathway (Fig. 5b). Initial investigations revealed that substrate **1a** exhibited negligible conversion under nitromethane-free conditions, with no detectable formation of the putative iminoquinone intermediate **16**, suggesting that **16** is not a stable off-pathway intermediate. To further assess its viability, chemically synthesized **16** was subjected to standard reaction conditions. While 14 efficiently produced 3a non-enzymatically in the presence of nitromethane, enzymatic catalysis yielded only trace amounts of **3a** (1% yield). The stark contrast between efficient non-enzymatic turnover and negligible enzymatic conversion of authentic **16** excludes **Pathway a** within the enzymatic mechanism. Additional evidence was obtained through high-resolution mass spectrometry (HRMS) analysis, which detected putative oxidative coupling intermediates **17** and **18**, derived from **1a** and methyl-phenol **1z**, respectively (Figure 5b; Supplementary Fig. 13 and 14). Furthermore, radical trapping experiments using butylated hydroxytoluene (BHT) severely suppressed the formation of **3a**, and the corresponding radical adduct **19** was successfully detected (Fig. 5c; Supplementary Fig. 15). Collectively, these results provide compelling support for the radical oxidative coupling pathway (**Pathway b**) as the predominant mechanism.

We further established the superior thermodynamic feasibility of **Pathway b** through DFT calculations. The mechanism initiates with a proton transfer from the phenolic hydroxyl group of substrates **1r** to the carboxylate of E502 via proton-coupled electron transfer (PCET) mechanism^55^, accompanied by concerted electron delocalization to form **INT1**. This initial step exhibits an activation barrier of 23.34 kcal/mol, thereby priming the substrate for subsequent transformation (Supplementary Fig. 16). Subsequently, a QM cluster model of **INT1’**, comprising **INT1**, nitromethane, and an OH^−^ group acting as a base, was used to model the formation of **INT3** (Fig. 5d). **Pathway a** involves three transition states, two of which exhibit high energy barriers: 26.59 kcal/mol for **TS2’** and 26.70 kcal/mol for **TS4’**. In contrast, **Pathway b** proceeds via radical oxidative coupling between the **INT1** and nitromethane with a significantly lower energy barrier of 13.94 kcal/mol (**TS2**). The resulting **INT2** is then enzymatically oxidized to form the thermodynamically stable **INT3**. Comparative analysis of the energy profiles confirms that **Pathway b** is energetically favored over **Pathway a** for the **INT1’**-to-**INT3** transformation, which is consistent with the preliminary experimental findings.

## Discussion

This study establishes a novel MCO-catalyzed system for skeletal editing, converting phenol and indole derivatives into value-added functionalized tropones and quinoline via exogenous single-carbon insertion in one step. This strategy employs nitroalkanes as stable, sustainable single-carbon synthons and O_2_ as the terminal oxidant, providing an advantageous alternative to traditional carbene-insertion protocols. Mechanistic studies integrating experimental and computational (DFT) approaches elucidate that the transformation proceeds through an MCO-mediated radical oxidative coupling between phenols and nitroalkanes to generate dienone, followed by a spontaneous tandem sequence of Michael addition and ring-expansion rearrangement. Engineered **BacFre** efficiently synthesizes structurally diverse tropone derivatives that exhibit superior antibacterial activity against to multidrug-resistant strains compared to their phenolic substrates. This strategy offers an operationally simple, mild, scalable, and environmentally benign platform to accelerate drug discovery through modular and atom-economy scaffold diversification.

## Methods

### Expression of multicopper oxidases

*E. coli* (*E. cloni* BL21(DE3)) cells harboring plasmids encoding the appropriate multicopper oxidases were culutred overnight in 5 mL of LB medium (kanamycin 50 µg/mL) at 37 °C with 250 rpm shaking. The pre-culture (0.5 mL) was inoculated into 20 mL of TB medium (kanamycin 50 µg/mL) in a 50 mL Erlenmeyer flask, followed by incubation at 37 °C with 220 rpm agitation. When the OD_600_ reached 0.6, 0.2 mM IPTG and 1.5 mM CuSO_4_ were added to the culture. Expression was conducted at 25 ºC with shaking at 180 rpm for 16–18 h. Following this, *E. coli* cells were pelleted by centrifugation (6000 × g, 10 min, 4 ºC). The media was removed, and the resulting cell pellet was resuspended in M9-N or MOPS buffer to an OD_600_ of 10 for the biotransformation.

### Purification of BacFre mutants

*E. coli* (*E. cloni* BL21(DE3)) cells harboring plasmids encoding **BacFre** mutants were cultured overnight in 6 mL LB medium (kanamycin 50 µg/mL) at 37 °C with 250 rpm shaking. The pre-culture (6 mL) was inoculated into 600 mL Terrific Broth (TB, kanamycin 50 µg/mL) in a 2-liter baffled flask, followed by incubation at 37 °C with 220 rpm agitation. When the OD_600_ reached 0.6, 0.2 mM IPTG and 1.5 mM CuSO_4_ were added to the culture. Expression was conducted at 25 ºC with shaking at 180 rpm for 16–18 h. Cultures were then centrifuged (6,000 × g, 20 min, 4 ºC) and the cell pellets were flash frozen with liquid nitrogen. For purification, frozen cells were resuspended in HisTrap buffer A (25 mM Tris, 100 mM NaCl, 20 mM imidazole, pH 7.5) and lysed by sonication (2 × 2 min, output control 5, 80% duty cycle). Lysates were centrifuged using a Lynx 6000 superspeed centrifuge (12,000 × g, 30 min, 4 ºC) to pellet cell debris. The protein was purified with a Ni-NTA column (5 mL HisTrap HP, Cat. No. 17524801, Cytiva) using an AKTA Start protein purification system. Proteins were eluted on a linear gradient from Histrap buffer A to His-trap buffer B (25 mM Tris, 500 mM imidazole, 100 mM NaCl, pH 7.5) about 12 column volumes (CV). Fractions containing the desired protein were collected, concentrated, and subjected to three exchanges of MOPS (50 mM, pH 8) using ultracentrifugal filters (20 kDa molecular weight cut-off, Amicon Ultra, Sigma Millipore) to remove excess salt and imidazole. Concentrated proteins were aliquoted with 20% glycerol, flash-frozen in liquid nitrogen and stored at -80 ºC until further use. The concentration of the protein sample is determined by BCA assay using Thermo Scientific’s protein BCA assay kit prior to use.

### General procedures for skeletal editing via single-carbon insertion using whole *E. coli* cells

#### (1) General procedures of reaction condition optimization

Suspensions of *E. coli* (*E. cloni* BL21(DE3)) cells expressing the corresponding multicopper oxidases in M9-N or MOPS buffer (typically OD_600_ = 10) and place it on ice for later use. To a 4 mL vial was added a suspension of *E. coli* cells expressing multicopper oxidases in prepared in buffer prepared in advance (350 µL), substrate **1a** (0.004 mmol, 40 µL of 0.1 M stock solution in MeCN), nitromethane **2a** (0.02 mmol, 10 µL of 2.0 M stock solution in MeCN) and H_2_O_2_ (1 µL, 30%). The vial was stirred at room temperature at 360 rpm for 12 h. Upon completion, the reaction was quenched with 800 µL MeCN containing 50 mM internal standard benzophenone. The mixture was stirred at room temperature at 500 rpm for 20 min and then transferred to a 1.5 mL Eppendorf tube and centrifuged (12,000 × g, 3 min). The supernatant was filtered through a 0.22 µm Millipore filter and subjected to reversed-phase liquid chromatography analysis to determine the reaction yield. Notes: reaction performed with *E. coli* cells resuspended to OD_600_ = 10 indicates that 350 µL of OD_600_ = 10 cells were added.

#### (2) General procedures of substrate scope exploration

To a 100 mL Erlenmeyer flask containing substrate **1** (0.2 mmol) and nitromethane **2** (2–6 mmol) were added MeCN (2 mL) and a suspension of *E. coli* cells expressing the **BacFre-K474F/I500L** variant in MOPS buffer (18 mL, typically OD_600_ = 10). The flask was shaken at room temperature at 220 rpm for 12–36 h. Upon completion, the reaction was extracted with 3 × 40 mL MeCN/EtOAc (4:1), and NaCl was added to promote the separation of the organic phase and the aqueous phase. The organic phases were combined, dried over anhydrous Na_2_SO_4_, filtered and concentrated in vacuo. The crude residue was purified by silica gel column chromatography to afford product **3**. Product **3** had poor solubility in ethyl acetate (EtOAc) and dichloromethane (DCM).

### Enzymatic reactions using purified protein

To a 4 mL vial were added MOPS buffer (268 µL), CuSO_4_ (0.00004 mmol, 4 µL of 0.01 M stock solution in ddH_2_O) and purified enzyme **BacFre-K474/I500L** (0.00004 mmol, 78 µL, 32.3 mg/mL in MOPS). After stirring for 10 min, substrate **1a** (0.004 mmol, 40 µL of 0.1 M stock solution in MeCN) and nitromethane **2a** (0.12 mmol, 10 µL of 12 M stock solution in MeCN) were added to the mixture. The vial was stirred at room temperature with 360 rpm for 12 h. Upon completion, the reaction was quenched with 800 µL MeCN containing 50 mM internal standard benzophenone. The mixture was stirred at room temperature at 500 rpm for 20 min and then transferred to a 1.5 mL Eppendorf tube and centrifuged (12,000 × g, 3 min). The supernatant was filtered through a 0.22 µm Millipore filter and subjected to reversed-phase liquid chromatography analysis to determine the reaction yield.

### Reporting summary

Further information on research design is available in the Nature Portfolio Reporting Summary linked to this article.

## Data availability

The experimental procedures and characterization data, NMR spectra of the compounds, mechanistic studies and computational calculations in this article are provided in the Supplemental Information. The structures of compounds **3a** (CCDC: 2481996), **4d** (CCDC: 2481997) in this article have been deposited in the Cambridge Crystallographic Data Center. These data could be obtained free of charge from The Cambridge Crystallographic Data Centre via www.ccdc.cam.ac.uk/data_request/cif

## Acknowledgments

This work was supported by the National Key R&D Program of China (2022YFA0912000), the Center of Synthetic Biology and Integrated Bioengineering (WU2022A006, WU2022A007, WU2023A009), the Research Center for Industries of the Future (WU2022C032). We thank Instrumentation and Service Center for Molecular Sciences (ISCMS) at Westlake University for the assistance in measurement and data interpretation. We thank Dr. Yinjuan Chen for the assistance with high resolution mass spectrometry (HRMS), Dr. Xiaohe Miao for the assistance with X-ray diffraction experiments.

## Author contributions

B.J. and Y.W. conceived and designed the overall project. B.J and J.-X.H. performed all synthetic experiments and wet mechanism experiments. H.-X.G, X.L., and M.-J.M. created mutations. B.-B.C. carried out the computational studies and Gibbs reaction energy (ΔG). B.J. and Y.W. wrote the paper with input of all authors.

## Additional information

## Supplementary Information

### Supplementary Date

### Competing interests

The authors declare no competing interests.

**Reprints and permission** information is available online at http://npg.nature.com/reprints and permission

### Publisher’s note

Springer Nature remains neutral with regard to jurisdictional claims in published maps and institutional affiliations.

